# Enhancing Lipid Detection and Spatial Accuracy in Carotid Plaques Using Mass Spectrometry Imaging Techniques

**DOI:** 10.64898/2026.01.14.699204

**Authors:** Sphamandla Ntshangase, Sean McCafferty, Zahra Javid, Beth Whittington, Phyo Khaing, Tony Feng, Allison Winarski, Fraser Simpson, Shazia Khan, Joanna P Simpson, Hannah O’Neill, Sharon Yen Ming Chan, Alaa Hassouneh, Mark Davis, Rahul Velineni, Andrew Tambyraja, Alexander von Kriegsheim, Catriona Graham, Rachael Forsythe, Stephanie Sellers, Patrick WF Hadoke, David E Newby, Ruth Andrew

**Affiliations:** University/BHF Centre for Cardiovascular Science, University of Edinburgh, Edinburgh, UK; Department of Vascular Surgery, Royal Infirmary Edinburgh, Edinburgh, UK; Cancer Research UK Scotland Centre, Institute of Genetics and Cancer, University of Edinburgh, Crewe Road South, Edinburgh, UK; Edinburgh Clinical Research Facility, The University of Edinburgh, Edinburgh, UK; Centre for Heart Lung Innovation, St Paul’s Hospital and University of British Columbia, Vancouver, Canada

## Abstract

Matrix-assisted laser desorption/ionisation mass spectrometry imaging (MALDI-MSI) is a powerful technique for studying lipid distribution in carotid plaques, key to understanding atherosclerosis. This study aimed to improve sample preparation for MALDI-MSI-based spatial lipidomics of carotid plaques by improving both matrix application and tissue handling.

Human carotid plaques were collected from endarterectomy patients with ethical approval and sectioned at 10 µm thickness for MALDI-MSI. We compared eight sample preparation methods, including hydroxypropyl methylcellulose-polyvinylpyrrolidone (HPMC-PVP) embedding media and Cryofilm-type IMS® to provide support and maintain tissue structural integrity during sectioning. Methods were assessed for signal intensity, lipid diffusion, lipid coverage, tissue morphology, and image co-registration which each criterion scored from 1-3.

Cryofilm-based methods scored highest for preserving tissue morphology and minimising folding artifacts (2.9-3.0) but were limited in co-registration (2.0) due to reliance on adjacent sections. Sublimation methods generally produced greater lipid coverage with reduced lateral diffusion, while automated sprayer methods scored higher in signal intensity/sensitivity (3.0) but had increased lipid delocalisation, particularly for highly hydrophobic species such as triacylglycerols and sterols.

The results highlight clear trade-offs between tissue structural preservation, lipid detection sensitivity, and spatial integrity in MALDI-MSI. Because spatial integrity cannot be compromised for imaging lipids in carotid atherosclerotic plaques, Cryofilm combined with sublimation offers a clear advantage. This work strengthens MALDI-MSI workflows enabling more precise spatial mapping and deeper biological interpretation of atherosclerotic lipid distributions.

## Introduction

Matrix-assisted laser desorption/ionisation (MALDI) mass spectrometry imaging (MSI) has become a transformative technique for mapping lipid distributions in biological tissues, especially in complex tissues like carotid plaques[1]. Such plaques contribute to arterial narrowing or atherosclerosis, a prevalent contributor to ischemic heart disease and stroke. Accurate lipid spatial imaging of plaques is key to advancing our understanding of their pathological development and this can be achieved by MALDI-MSI. However, lipid delocalisation remains a challenge in MALDI-MSI, as the native spatial distribution of certain lipid classes in tissues can be distorted. This undermines imaging accuracy and hampers the localisation of lipids to their biological origins, potentially leading to misinterpretations. Sensitivity of detection is another concern as delocalisation of some lipid species may lead to ion suppression of others [2]. Low-abundance lipids are particularly susceptible, as their signals may fall below the detection threshold due to inadequate extraction or ion suppression. Ensuring robust sensitivity across a wide spectrum of lipids is essential for comprehensive lipid profiling.

Matrix application methods play a pivotal role in addressing this challenge. Conventionally, automatic sprayers offer precise control over matrix deposition e.g., through adjustable gas flow and nozzle velocity [3]. Although this solvent-based approach improves matrix co-crystallisation with analytes, it often exacerbates lipid delocalisation, complicating accurate lipid localisation in tissues [4]. Sublimation, a widely used technique involving the controlled heating of matrix crystals, can yield a fine, uniform coating that enhances spatial resolution, with crystal sizes less than 1 μm [5, 6]. Despite its strengths, sublimation may result in limited lipid extraction due to lack of solvent, potentially reducing signal intensity. Thus, challenges lie in achieving effective matrix-analyte interaction without causing delocalisation, which may be addressed by carefully introducing solvent/vapour to induce matrix-analyte recrystallisation [7].

Different tissue preparation methods have also been investigated to improve localisation. For example, hydroxypropyl methylcellulose-polyvinylpyrrolidone (HPMC-PVP) embedding has been used to preserve tissue integrity [8]. However, some embedding media have been reported to introduce interferences and even cause delocalisation [9, 10]. Cryofilm, a polymer film onto which tissue sections are applied, has been shown to maintain lipid spatial integrity, particularly in challenging heterogeneous and brittle tissues like bones [11].

This study systematically evaluated the potential benefits of sublimation *vs* automated spray-based matrix deposition methods and the relative value of the different tissue mounting techniques (HPMC-PVP embedding and Cryofilm type IMS®) to assess their impact on lipid localisation and sensitivity of detection in carotid plaques. By systematically comparing these approaches, the research aims to develop optimised workflows that minimise lipid delocalisation while preserving the spatial accuracy and detection thresholds required for effective MALDI imaging.

## Experimental Section

### Tissue Processing

Carotid plaque specimens were obtained with informed consent from male patients undergoing carotid endarterectomy at The Royal Infirmary of Edinburgh, NHS Lothian, in accordance with ethical regulations. The tissue was cut into 10 μm serial cryosections using an MX35 Premier Blade (Fisher Scientific, UK) and a CryoStar NX50 cryostat (Fisher Scientific) at −16 °C. Sections were thaw-mounted either directly onto SuperFrost Plus slides (Fisher Scientific, UK) or Cryofilm type IMS® (SECTION-LAB, Hiroshima, Japan). Following sectioning, the Cryofilm type IMS® with tissue was adhered to a slide prepared with copper strip (SECTION-LAB Co. Ltd.) and ZIG 2-way glue (Kuretake Co., Ltd., Nara, Japan) as shown in Figure 3.

### Matrix Application

An automated spraying method previously reported by Ntshangase et.al[12] was used in the first instance. We contrasted this to a sublimation approach using an HTX Sublimator system (HTX-Technologies, Chapel Hill, North Carolina, USA). For the latter, the α-cyano-4-hydroxycinnamic acid (CHCA) (Sigma-Aldrich Gillingham, UK) matrix solution (50 mg/mL; 2 mL) was prepared in 70/29.9/0.1 acetonitrile (LC-MS grade, Fisher Scientific, Loughborough, UK)/water (LC-MS grade, Fisher Scientific, Loughborough, UK)/trifluoroacetic acid (Sigma-Aldrich Gillingham, UK), and poured (2 mL) onto the sublimation plate. The solvent was evaporated prior to sublimation by setting the plate temperature at 70 °C. The CHCA matrix was applied onto the tissue sections by sublimation (180 °C, 5 min, vacuum starting at <0.004 mBar). Subsequently, and when desired, the matrix was recrystallised in an oven at 60 °C using preheated 400 µL 50% methanol (v/v) (LC-MS grade, Fisher Scientific, Loughborough, UK)/0.1% trifluoroacetic acid in 1 L sealed plastic box for 30 seconds.

### MALDI Mass Spectrometry Imaging

MALDI MS data were acquired in positive mode (*m/z* 300–1000) on a Synapt G2Si MS TOF mass spectrometer (Waters, Billerica, MA) using a 355 nm Nd:YAG laser at kHz frequency. External calibration was performed with red phosphorus signals prior to data acquisition. MSI was conducted in sensitivity mode with 150 mJ laser energy and a spatial resolution of 75 × 75 μm^2^, acquiring 1 scan/0.5 seconds. Trap and transfer collision energies were set to 4 V and V, respectively, yielding a mass resolution of approximately 17,000 full-width at half-maximum (FWHM) at *m/z* 400.

### Histology

Following MS imaging, the MALDI matrix was removed with 50%, 70% then 100% ethanol (v/v) washes for 20 sec each time. The samples were stained with hematoxylin and eosin (H&E) and cover slipped with Permount (Thermo Fisher Scientific, UK). An H&E image was generated at 20x magnification using an Odyssey M Imaging System (Li-COR, Biosciences, UK).

### Data Processing

Raw MSI data were processed using LipostarMSI (Molecular Horizon Srl, Perugia, Italy). Lock-mass correction of averaged spectra was applied using sphingomyelin SM 34:1 detected as [M+Na]^+^. Mass filtering was performed using tolerances of 0.05 Da for Synapt data and 0.0025 Da for FT-ICR data. Ion images underwent hotspot removal using a 99.98% threshold. A minimum ion intensity threshold was applied, and any peaks below this threshold were discarded. Whole-tissue regions of interest (ROIs) were manually defined based on corresponding histological sections and optical images, with the vessel lumen excluded to ensure accurate tissue representation. ROI spectra were normalised to total ion current (TIC), and the resulting data matrices were exported for downstream analysis, visualisation using Python (version 3.12.4) and GraphPad Prism (version 10.2.3).

### Lipid Annotation

Lipids were subsequently annotated by acquiring further spectra on a 12T solariX MALDI-FT-ICR-MS (Bruker Daltonics) using ftmsControl v2.2 and matching data to the LIPID MAPS database with <5 ppm mass tolerance. Spectra were acquired. A SmartBeam-II laser (355 nm) operated at 1 kHz, 200 shots per pixel and 40% power in medium focus mode (75 × 75 μm^2^ ablation area). Data were collected across *m/z* 100–1200 with an estimated mass resolution of ∼130,000 at *m/z* 400, and processed in LipostarMSI. Selected ions were fragmented on-tissue using MALDI MS/MS (Synapt G2-Si) with varying collision energy (20–40 V) and low mass (LM) resolution was set at 15.

### Scoring System

To evaluate and compare sample preparation methods for spatial lipidomics, we developed a scoring system based on key performance indicators. Each method was assessed against five criteria including lipid signal intensity, diffusion, coverage, tissue folding artifacts, and image coregistration. Table 1 outlines the scoring system used.

**Table 1.**
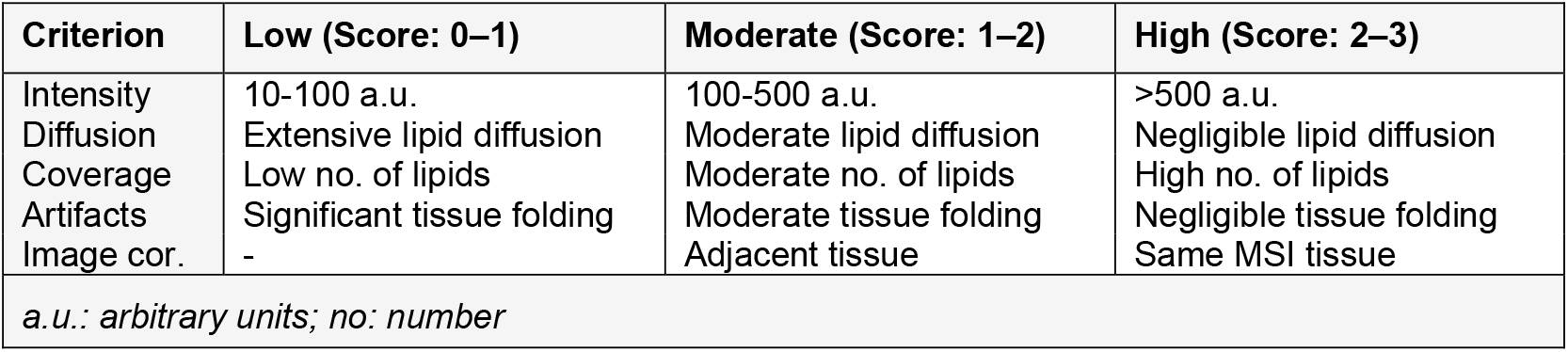
Scoring criteria for evaluating lipidomic sample preparation methods.

## Results and Discussion

### Tissue Preparation for MALDI Mass Spectrometry Imaging

Optimising sample preparation is essential for reliable and reproducible spatial lipidomics by MALDI-MSI. Solvent-based matrix application methods, while commonly used [12], are inherently limited by solvent-induced lipid delocalisation, which compromises spatial integrity and increases the risk of false biological interpretations [9]. This problem is particularly evident in lipid-rich tissues such as human carotid plaques, where maintaining the native spatial distribution of lipids is important for accurate molecular mapping. To address this, we evaluated eight tissue preparation methods (Table 2) combining different tissue supports (none, HPMC-PVP, Cryofilm, or both) with CHCA matrix deposition using either an automated sprayer (supplementary information)[12] or a sublimator (with or without recrystallisation). While matrix spraying remains the most widely used approach, sublimation is being increasingly adopted in high-resolution lipid imaging due to its improved spatial precision [13].

**Table 2.**
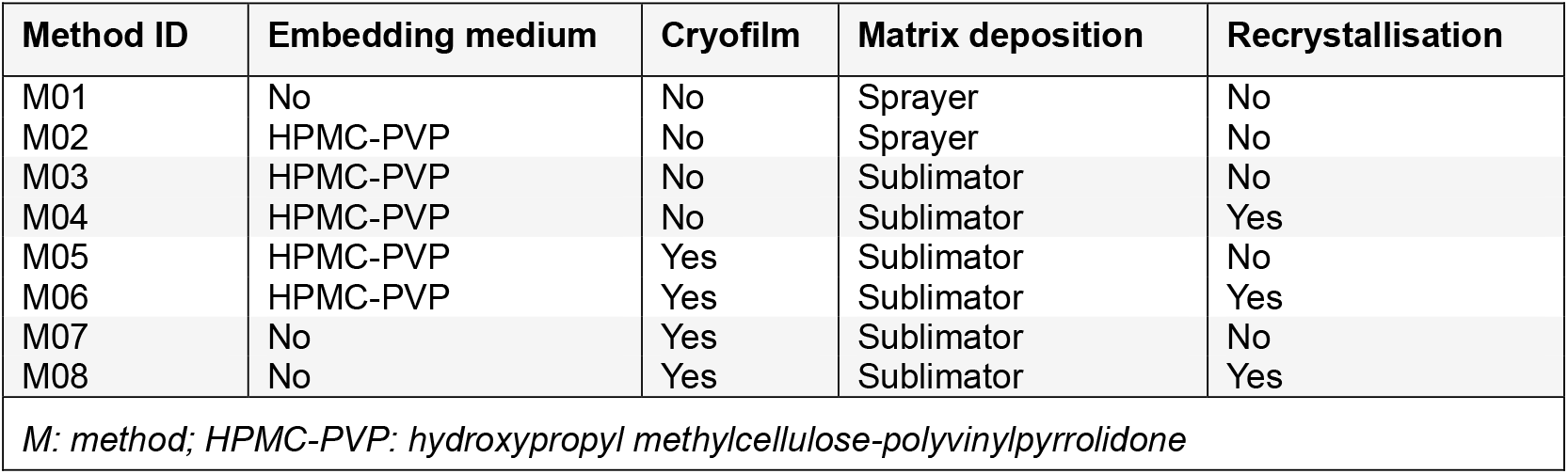
Tissue preparation methods differing in embedding, Cryofilm use, matrix deposition and recrystallisation.

### Initial Screening

Early experiments using conventional solvent-spray methods (M01) showed evident tissue folding that compromised structural integrity, clearly visible to the naked eye (Figure S2). However, as expected, embedding the tissue in HPMC-PVP (M02) moderately restored the tissue structure (Figure S3) [12]. Despite the strong signal intensity observed in both methods, there was evident lipid diffusion, with exemplar lipids including lysophosphatidylcholines (e.g., [LPC 16:0 + Na]^+^, *m/z* 518.32), phosphatidylcholines ([PC 34:1 + Na], *m/z* 782.57), and loss of signal for sterols (e.g., [ST 27:1;O+H - H_2_O]^+^, *m/z* 369.35) and triacylglycerols (TGs) (Figure 1). We observed that lipid classes like sterols and triacylglycerols frequently delocalised during solvent-based methods, forming droplets across the tissue and accumulating at or beyond the tissue edges (Figure S3). Their extreme hydrophobicity causes them to migrate with the solvent, leading to spreading, loss of spatial integrity, and reduced signal intensity.

**Figure 1.**
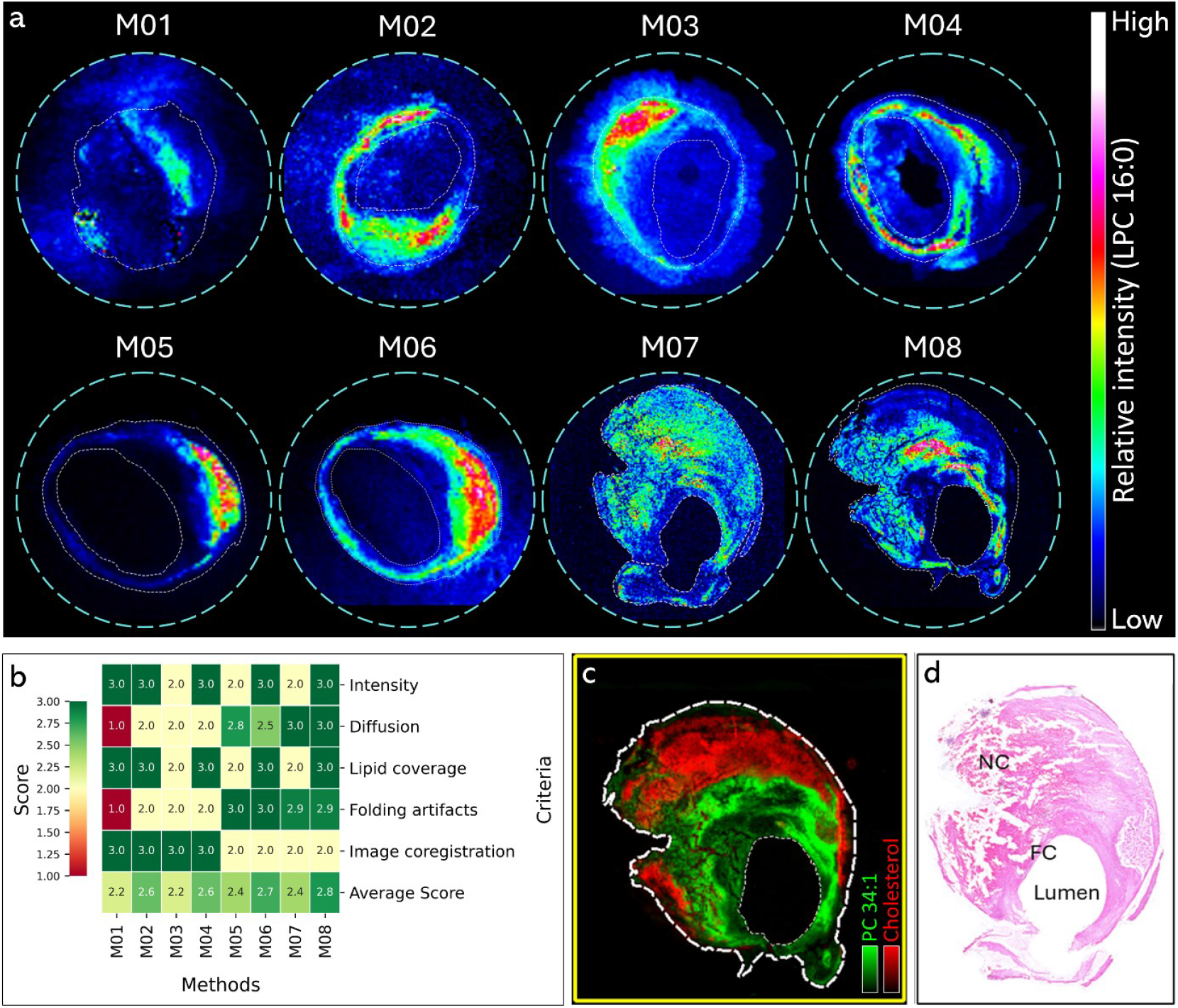
**a)** Exemplar MALDI MS images of [LPC 16:0 + Na]^+^ at *m/z* 518.32 comparing eight sample preparation methods (M01-M08). **b)** The scoring criteria of the performance of eight tissue preparation methods based on lipid intensity, diffusion, coverage, tissue folding artifacts and image coregistration. Eight sample preparation methods were assessed including applying the matrix with an automated sprayer (M01-M02) or sublimation (M03-M08). Each method was scored on a 0–3 scale, where higher scores indicate better performance. Tissue embedded in HPMC-PVP + Cryofilm + recrystallisation (M06) and sectioning tissue directly onto the Cryofilm + recrystallisation (M08) showed the strongest performance (average score = 2.7 and 2.8 respectively). **c)** MALDI-MSI images (M08) showing phosphatidylcholine, [PC 34:1+ Na]^+^ at *m/z* 782.57 localised in fibrous tissue and cholesterol [ST 27:1;O + H-H_2_O]^+^ at *m/z* 369.35, mostly in the necrotic core. **d)** Adjacent H&E-stained section highlighting the fibrous cap (FC) and necrotic core (NC) regions.

In contrast, some sublimation-based methods produced sharper boundaries for most lipids and improved spatial integrity (Figure 1a). We evaluated six sublimation-based methods (M03-08) in search of enhanced spatial integrity for lipid imaging in carotid plaques. The first sublimation method (M03) involved initial embedding of tissue in HPMC-PVP followed by matrix application *via* sublimation. This method slightly improved localisation of extremely hydrophobic lipids like sterols (e.g., cholesterol and derivatives) but its poor matrix co-crystallisation reduced lipid extraction efficiency and signal sensitivity of most lipid species across the spectrum (Figure. 1a). For some lipids, delocalisation persisted even without spraying solvent into the surface of the tissue (e.g., lysophosphatidylcholines), suggesting the influence of additional factors including the embedding medium, slide surface properties, and temperature effects. Introducing matrix recrystallisation (M04) improved detection and intensity (Figure. 1a) but did not fully resolve concerns about delocalisation.

Applying Cryofilm type IMS® substantially improved tissue structure (M05-08), yet diffusion remained without (M05) or with (M06) recrystallisation, largely due to residual water content from HPMC-PVP hydrogel, which particularly affected highly hydrophobic lipid classes like sterols and triacylglycerols. During optimisation, we discovered that the water content in HPMC-PVP caused the hydrogel to rapidly disintegrate upon contact with the hydrophobic Cryofilm surface, leading to partial lipid delocalisation at the tissue edges. We thus explored the Cryofilm alone as a mechanical support to eliminate the need for HPMC-PVP. Indeed, omitting the embedding media entirely prevented delocalisation while tissue morphology remained preserved. Direct mounting of tissue onto Cryofilm type IMS® without HPMC-PVP (M07-08), combined with recrystallisation (M08), preserved sensitivity comparable to HPMC-PVP with recrystallisation (M04) while effectively eliminating diffusion and delocalisation.

### Systematic Scoring

The scoring criteria based on signal intensity collated across the whole tissue, lipid diffusion, lipid coverage, tissue folding artifacts, and image coregistration (Table 1) showed clear trade-offs between the eight methods. Following sublimation, methods with recrystallisation achieved the highest scores for intensity (score = 3.0) and lipid coverage (score = 3.0) (Figure 1b). HPMC-PVP + Cryofilm-assisted methods with recrystallisation maximised spatial retention (score = 2.5 - 3.0) while maintaining higher intensity and lipid coverage. HPMC-PVP without recrystallisation showed moderate diffusion (score = 2.5) but lower lipid coverage (score ≤ 2.0), and HPMC-PVP + Cryofilm without recrystallisation scored lower in intensity (score = 2.0) and lipid coverage (score = 2.2) despite excellent spatial retention (score = 2.8). Cryofilm without HPMC-PVP but paired with recrystallisation (M08) achieved the highest lipid coverage (score = 3.0) with negligible to non-existent diffusion (score = 3.0).

Regarding tissue morphology, all Cryofilm-assisted methods performed best in maintaining tissue integrity and minimising folding artifacts introduced during sectioning (scores = 2.9-3.0). However, they scored lower in image co-registration (score ≤ 2.0) because these methods rely on adjacent sections rather than the same tissue, illustrating the inherent limitation that a single section cannot be used for both histology and MS imaging since the Cryofilm is not transparent.

### Highest Scoring Methods - A Direct Comparison

Following iterative assessments of different approaches, we identified two top-performing sublimation-based methods, HPMC-PVP + Cryofilm + recrystallisation (M06) and Cryofilm + recrystallisation (M08). These two methods were selected for direct comparison, as both represent refined approaches that balanced lipid spatial preservation and detection sensitivity. The M08 approach frequently generated high-intensity signals of the common abundant ions (e.g., a fold change of 1.32 for cholesterol and 1.95 for PC 34:1), whereas the M06 method yielded a broader distribution of intensities with relatively fewer dominant peaks (Figure 2a). The distinct spectral profiles imply that each preparation method enhances or suppresses the detection of different ion populations. However, shared signals were observed in certain *m/z* intervals, suggesting negligible effect on some lipids.

**Figure 2.**
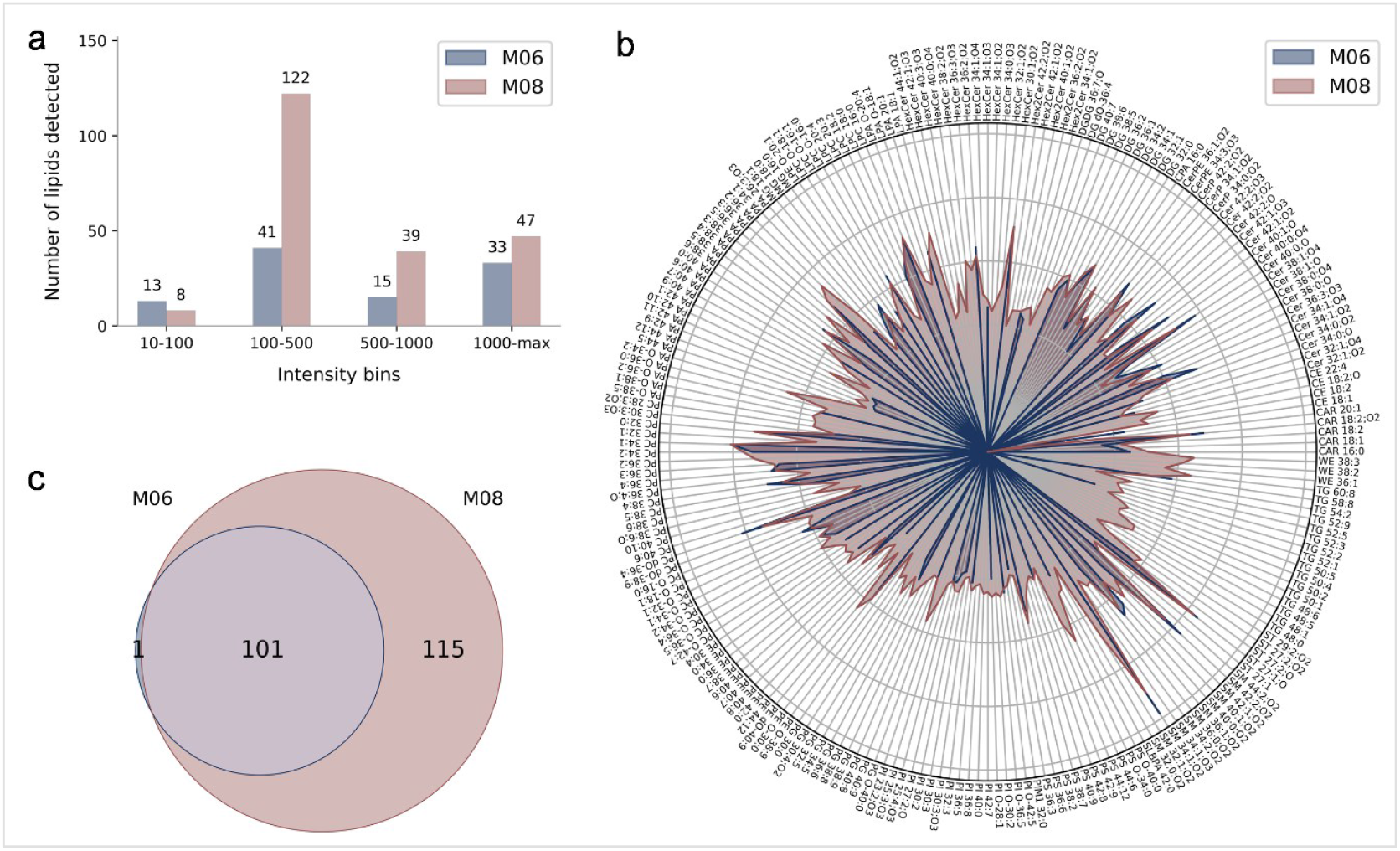
Comparative lipid coverage of the two best sample preparation methods from initial screening including HPMC-PVP + Cryofilm-recrystalised (M06) and Cryofilm-recrystalised (M08). (**a**) Bar graph depicting the distribution of lipid subclasses detected in HPMC-PVP + Cryofilm-recrystalised (M06) and Cryofilm-recrystalised (M08) across different intensity thresholds. (**b**) Radar chart illustrating the relative abundance of individual lipid species detected in HPMC-PVP + Cryofilm-recrystalised (M06) and Cryofilm-recrystalised (M08). Notable increases in lipid coverage in Cryofilm-recrystalised (M08) are observed across multiple classes, particularly within triacylglycerols (TGs) and hexosylceramides (HexCer). (**c**) Venn diagram showing the overlap of lipid species identified in HPMC-PVP + Cryofilm-recrystalised (M06) (102) and Cryofilm-recrystalised (M08) (216), with 101 common lipid species between the two methods.

To assess the lipidomic coverage between the two methods we compared the number of identified lipids across different intensity thresholds (Figure. 2a). A total of 102 lipid species were detected following HPMC-PVP + Cryofilm-recrystalised (M06), while 216 were identified after Cryofilm-recrystalised (M08), with 101 lipids common in both methods (Figure. 2c). A comprehensive radar chart visualisation revealed a higher coverage of several lipid classes in tissue sectioned directly onto the Cryofilm-recrystalised (M08) compared to HPMC-PVP + Cryofilm-recrystalised (M06), including triacylglycerols and hexosylceramides (HexCer). Diacylglycerols (DGs) remained minimally affected with a few detected only in Cryofilm-recrystallised method (M08). Some ceramides were only detected in the Cryofilm-recrystallised method (M08) (Figure. 2b). This suggests overall enhanced lipid diversity and accumulation in the datasets when applying the Cryofilm-recrystallised method (M08). Notably, certain lipid species, such as CE 18:1 and TG 48:5, showed substantial increase in Cryofilm-recrystallised method (M08), while sphingomyelins (SM) and phosphatidylcholine (PC) appeared more heterogeneous with some lost in M06.

### Optimised workflow

The conventional spray-based workflow is presented in Figure S1, and Figure 3 illustrates the optimised Cryofilm-assisted sublimator-based workflow (M08), specific for MALDI-MSI of lipids in carotid plaque tissue. It should be noted that this method works well for human carotid plaque tissue, which is relatively large (0.5-1 cm wide samples), but adapting it to very small tissues may present new challenges. HPMC-PVP + Cryofilm-recrystalised (M06) may be more suitable for such cases.

**Figure 3.**
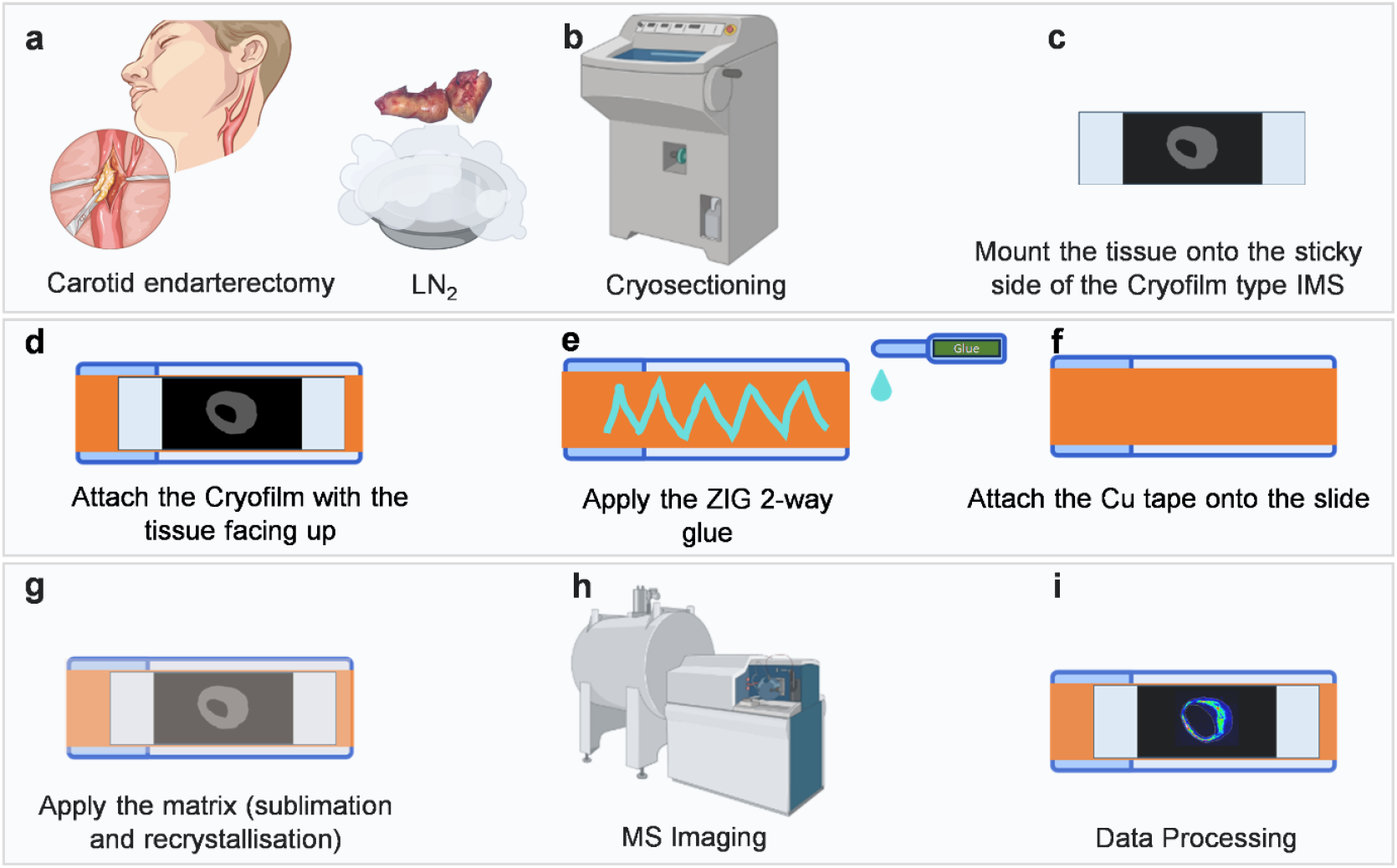
Workflow for Cryofilm type IMS®-assisted mass spectrometry (MS) imaging of human carotid atherosclerotic plaque. (a) Carotid endarterectomy tissue was collected from patients undergoing surgery. (b) The excised plaques were cryosectioned using a cryostat. (c) Tissue sections were collected on Cryofilm type IMS®, which preserves morphology and facilitates downstream handling. (d) The Cryofilm type IMS® with the tissue side facing up was transferred onto a glass slide. (e) by applying the ZIG 2-way glue (**NB** it was found important to wait until the glue turns clear before mounting the Cryofilm type IMS®, otherwise it would not dry). (f) The Cryofilm type IMS® adhesive tape was mounted securely onto the slide. (g) Matrix was applied by sublimation followed by recrystallisation to enhance analyte extraction and desorption/ionisation efficiency. (h) MS imaging was performed to acquire spatially resolved molecular data. (i) The acquired datasets are processed and analysed to visualise and interpret the spatial distribution of lipids or other molecular species across the tissue section.

## Conclusion

This study highlights key trade-offs in lipid coverage and localisation across eight different matrix application and tissue preparation methods. Cryofilm effectively eliminates delocalisation but at the cost of image coregistration. Sublimation without recrystallisation maintains spatial accuracy but reduces overall sensitivity. Importantly, solvent-driven delocalisation are major concerns that must be addressed for improved MALDI lipid imaging accuracy. Researchers should select their chosen approach depending on the specific tissue and biological questions under study.

## Acknowledgments

This work is funded by Heart Research UK (NET23-100005). For the purpose of open access, the authors have applied a Creative Commons Attribution (CC BY) license to any Author Accepted Manuscript version arising from this submission. Experiments were conducted at the Edinburgh Clinical Research MS Core (RRID:SCR_021833) and the Scottish Instrumentation and Research Centre for Advanced MS.

## Data Availability

All data will be uploaded to Zenodo. Additional data are available from the corresponding author on reasonable request.

## Notes

### Competing Interest Statement

The authors have declared no competing interest.

