## supplementary information for "Enhancing Lipid Detection and Spatial Accuracy in Carotid Plaques Using Mass Spectrometry Imaging Techniques"

### Methods

**Automated Sprayer Matrix Application:** Slides were coated with CHCA (10 mg/mL in 70/29.9/0.1 acetonitrile/water/TFA) using an HTX M3 sprayer (Table S1). Spray parameters for CHCA were as follows: temperature, 75 °C; flow rate, 0.24 mL/min; nozzle velocity, 1200 mm/min; track spacing, 1.0 mm; nitrogen pressure, 10 psi; drying time, 5 s; number of passes, 2; and matrix density, 400 μg/cm². The slides were subsequently scanned using an Epson Perfection V370 optical scanner (London, UK).

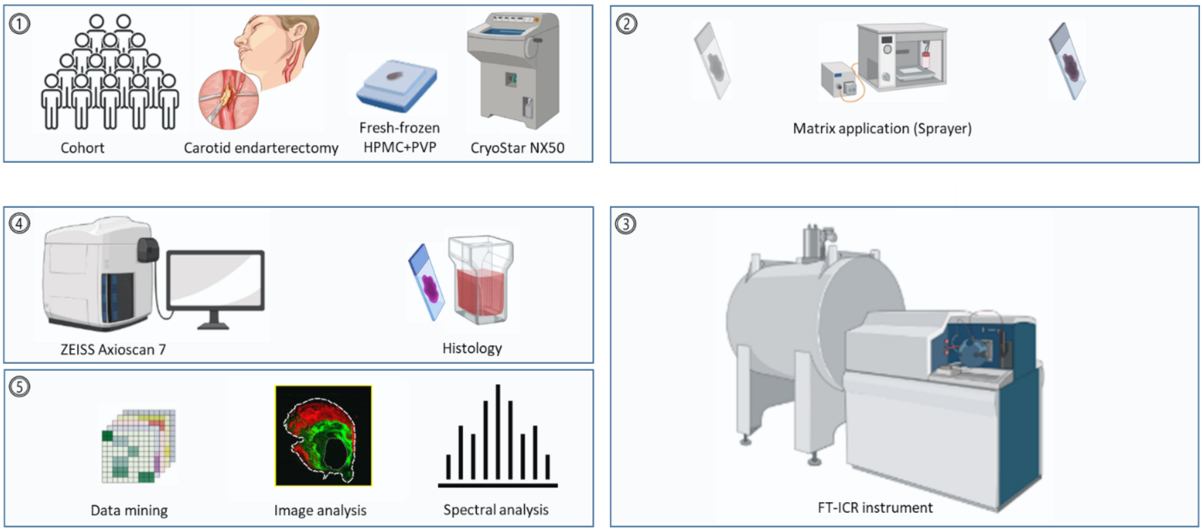


**Figure S1**. a) Overview of the experimental and analytical workflow. 1) Carotid endarterectomy specimens were collected from the patient cohort, fresh-frozen, and cryosectioned. Tissue sections were prepared for multimodal analysis, including 2) matrix application for 3) mass spectrometry imaging, FT-ICR MS acquisition, 4) histological staining, and optical imaging. 5) High-resolution imaging and spectral data were subsequently processed through image analysis, spectral analysis, and data mining to enable integrated spatial and molecular characterisation of carotid plaque tissue.

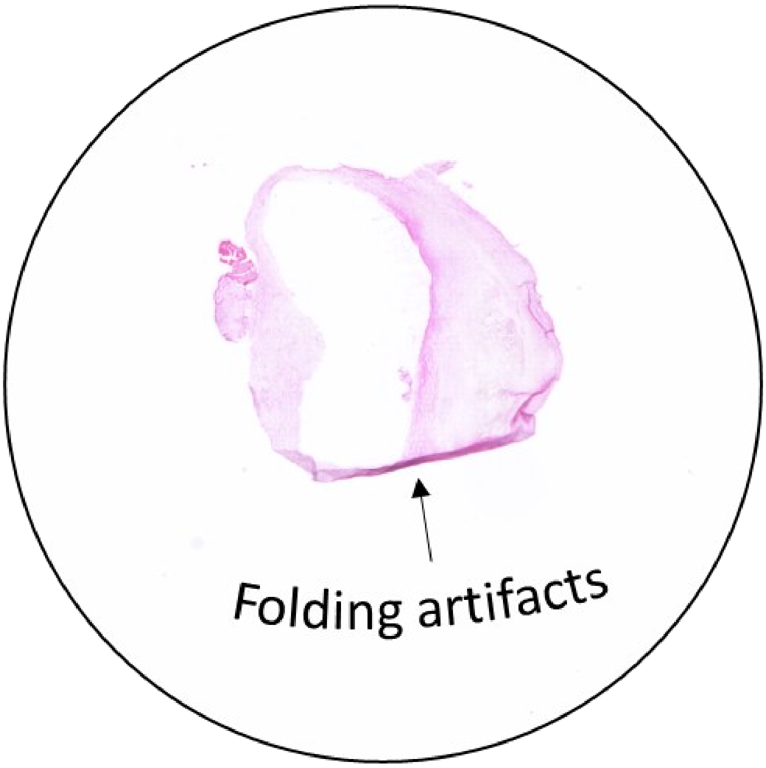


**Figure S2.** a) H&E image of a carotid plaque cryosection without supporting embedding media, showing visible folding artifacts.

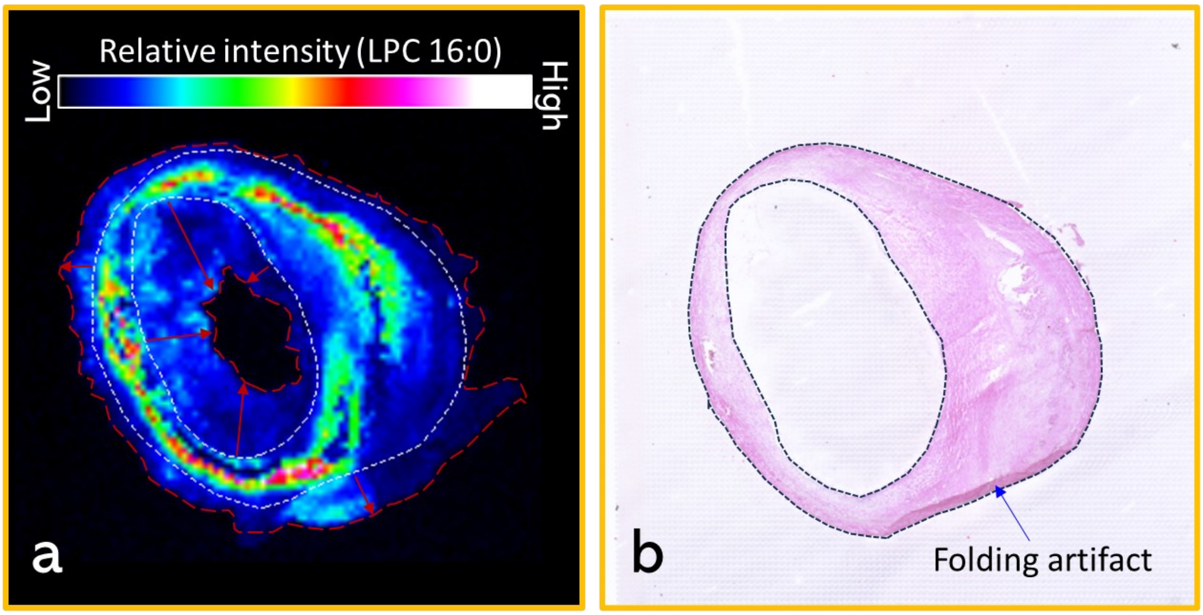


**Figure S3**. a) MALDI-MS image of [LPC 16:0 + Na]$^+$ at *m/z* 518.32 showing diffusion beyond the tissue. The white dotted line indicates the tissue boundary, while the red dotted line marks the extent of the diffusion. b) Hematoxylin and Eosin (H&E) staining of the same tissue analysed by MSI, with minor tissue folding artifacts.

**Table S1**. Tissue preparation methods differing in embedding, Cryofilm use, and matrix deposition – **matrix density (µg/cm²)**.

| Method ID | Embedding medium | Cryofilm type IMS | Matrix deposition | Recrystalised (Yes/No) | Matrix density (µg/cm²) |
|---|---|---|---|---|---|
| M01 | None | No | Sprayer | No | 400 |
| M02 | HPMC–PVP | No | Sprayer | No | 400 |
| M03 | HPMC–PVP | No | Sublimator | No | 160 |
| M04 | HPMC–PVP | No | Sublimator | Yes | 160 |
| M05 | HPMC–PVP | Yes | Sublimator | No | 160 |
| M06 | HPMC–PVP | Yes | Sublimator | Yes | 160 |
| M07 | None | Yes | Sublimator | No | 160 |
| M08 | None | Yes | Sublimator | Yes | 160 |

Matrix density (µg/cm$^2$) was calculated by the equation:

$$Density = \frac{Wa - Wb}{A}$$

$W_b$ = Weight of the clean glass slide or sample holder (in µg).
$W_a$ = Weight of the slide after the sublimation process is complete (in µg)
A = The total surface area of the slide is 18.75 cm$^2$ (in cm$^2$)
